# Precise prediction of antibiotic resistance in *Escherichia coli* from full genome sequences

**DOI:** 10.1101/338194

**Authors:** Danesh Moradigaravand, Martin Palm, Anne Farewell, Ville Mustonen, Jonas Warringer, Leopold Parts

## Abstract

The emergence of microbial antibiotic resistance is a global health threat. In clinical settings, the key to controlling spread of resistant strains is accurate and rapid detection. As traditional culture-based methods are time consuming, genetic approaches have recently been developed for this task. The diagnosis is typically made by measuring a few known determinants previously identified from whole genome sequencing, and thus is restricted to existing information on biological mechanisms. To overcome this limitation, we employed machine learning models to predict resistance to 11 compounds across four classes of antibiotics from existing and novel whole genome sequences of 1936 *E. coli* strains. We considered a range of methods, and examined population structure, isolation year, gene content, and polymorphism information as predictors. Gradient boosted decision trees consistently outperformed alternative models with an average F1 score of 0.88 on held-out data (range 0.66-0.96). While the best models most frequently employed all inputs, an average F1 score of 0.73 could be obtained using population structure information alone. Single nucleotide variation data were less useful, and failed to improve prediction for ten out of 11 antibiotics. These results demonstrate that antibiotic resistance in *E. coli* can be accurately predicted from whole genome sequences without *a priori* knowledge of mechanisms, and that both genomic and epidemiological data are informative. This paves way to integrating machine learning approaches into diagnostic tools in the clinic.

**Summary:** One of the major health threats of 21st century is emergence of antibiotic resistance. To manage its economic impact, efforts are made to develop novel diagnostic tools that rapidly detect resistant strains in clinical settings. In our study, we employed a range machine learning tools to predict antibiotic resistance from whole genome sequencing data for *E. coli.* We used the presence or absence of genes, population structure and isolation year of isolates as predictors, and could attain average precision of 0.93 and recall of 0.83, without prior knowledge about the causal mechanisms. These results demonstrate the potential application of machine learning methods as a diagnostic tool in healthcare settings.

## Introduction

Antibiotic resistance has turned into an acute global threat. The rise of bacterial strains resistant to multiple antibiotics is expected to dramatically limit treatment effectiveness [1], leading to potentially incurable outbreaks. In addition to new drug development efforts, there is an urgent need for preclinical tools that are capable of effective and rapid detection of resistance [2, 3], as culture-based laboratory diagnostics test are usually time consuming and costly [3].

To accelerate the diagnosis, genetic tests have been devised to identify known resistance genes. The increasingly affordable and available whole genome sequencing data from clinical strains has helped to robustly identify antibiotic resistance determinants, and to curate them in dedicated databases [4, 5]. Given sequence from a new strain, computational methods can then look up known causal genes in these resources [5, 6]. Whilst such rule-based models are highly accurate for some common pathogens with well-characterized resistance mechanisms (e.g. *Mycobacterium tuberculosis* and *Staphylococcus aureus*), they cannot be employed to detect resistance caused by unknown mechanisms in other major pathogenic strains, and require constant curation to remain effective.

Prediction approaches based on machine learning have the potential to overcome these restrictions of rule-based tests. As general-purpose methods, they are agnostic to the causal mechanisms, and learn useful features directly from data [7-9]. Already, decision tree based models have proven valuable for predicting resistance and pathogen invasiveness from genomic sequences [10-14]. However, these studies were limited in both the genetic features used and the methods applied. In particular, both population structure and accessory genome content could contain predictive information, as resistance determinants may be transferred horizontally from other strains, or inherited vertically from an ancestor [2]. Further, the powerful deep learning methods that can utilize complex features interactions were not examined.

Here, we systematically evaluate the performance of machine learning algorithms for predicting antibiotic resistance from *E. coli* whole genome sequence data. We present genome sequences and resistance measurements of 255 new isolates and consider them together with published data from recent large-scale studies, as well as simulated datasets. We test whether prediction accuracy improves with including temporal data, population structure, and accessory genome content, and assess how a range of population parameters, such as mutation and recombination rates, influence predictions.

## Methods

### Isolates

We used 1681 strains from four large-scale clinical and environmental *E. coli* collections, with available data on the year of isolation, drug susceptibility phenotypes, and whole genome sequence [15, 16]. Furthermore, we collected 255 strains from a range of ecological niches: hospital sewage and water treatment plant from Sweden (Carl-Fredrik Flach); human clinical isolates isolated in Pakistan, Syria, Sweden and USA (Culture Collection University of Gothenburg); a collection of strains producing extended-spectrum β-lactamases isolated in Sweden (Christina Åhrén) and environmental samples from Belgium (Jan Michiels).

### Antimicrobial susceptibility testing

Antimicrobials tested included beta-lactams (penicillin: ampicillin (AMP, 6µg/ml); cephalosporins: cefuroxime (CXM, 8µg/ml), cefotaxime (CTX, 4µg/ml), cephalothin (CET, 20µg/ml) and ceftazidime (CTZ, 0.25µg/ml)), aminoglycosides (gentamicin (GEN, 4µg/ml) and tobramycin (TBM, 8µg/ml)), and fluoroquinolones (ciprofloxacin (CIP, 1µg/ml)). Concentrations used were determined by performing a 2-fold serial dilution, starting from twice the concentrations listed by the European Committee on Antimicrobial Susceptibility Testing (EUCAST) on 25/01/2017, until no growth was observed after 16 hours for the common lab strain BW25113 [17] used as a control in the experiments.

### Sequencing data generation

We extracted DNA with the Bacterial Genomic DNA Isolation 96-Well Kit (Norgen Biotek) as detailed in the manufacturer’s instructions. Libraries were prepared with standard Illumina DNA sequencing library preparation protocols, and sequenced on Illumina HiSeq X with 150 bp paired end reads, multiplexing 384 samples per lane, and achieving average depth of coverage of 40-fold. We used Kraken, which accurately assigns taxonomic labels to the short DNA reads [18], to confirm the presence of *E. coli* reads in the pool. The raw sequences for the sequenced data in this study have been deposited in the European Nucleotide Archive (ENA) under the accession numbers described in Supplemental Table S1. The assembled data is available in the repository (www.github.com/DaneshMoradigaravand/PanPred).

### Pan-genome determination

Paired-end reads for the isolates sequenced both here and previously were assembled with Velvet [19] and put through an improvement pipeline [20]. In order to reconstruct the pan-genome, we used the output assemblies and annotated these with Prokka [21]. The annotated assemblies produced by Prokka were then used as input for Roary [22] to build the pan-genomes with the identity cut-off of 95%. Roary produced a matrix for the presence and absence of accessory genes. The variant sites (SNPs) in the core genome alignment were extracted with an in-house snp_sites tool (www.github.com/sanger-pathogens/snp-sites). To visualize the phylogenetic tree with the associated metadata, we used iTOL [23].

### Population structure calculation

We mapped the short reads to the reference EC958 genome sequence [24] as detailed in [25], and calculated the pairwise SNP distance (number of differing sites) for the core genome alignment of strains with functions in the ape package [26]. We identified clusters within the population using a distance-based method in the adegenet package [27]. We clustered sequences using the sequence distance metric with the adegenet package for all possible number of clusters from 1 to number of strains. Based on these clusterings, we constructed the population structure matrix S, where s_ij_ = *k* if strain *i* belongs to cluster *k* in the clustering with at most *j* clusters.

### Simulated datasets

To evaluate the performance of prediction tools, we simulated pan-genomes with Scoary [28]. The simulation process begins with a single genome with 3000 core and 6000 accessory genes that undergoes duplication and gene loss/gain in every generation, and continues until a desired number of genomes is reached; we tested population sizes of 130, 260, 650 and 1300. We examined penetrances, defined as the probability of acquisition/loss of the resistance phenotype simultaneously to the acquisition/loss of the causal resistance gene, of 0.5, 0.6, 0.7, 0.8, 0.9 and 1.

### Feature calculation

We examined different predictors as inputs: 1) matrix of the presence-absence of accessory genomes within the pan-genome (G), where g_ij_ is 1 if gene *i* is present in strain *j*, and 0 otherwise; 2) matrix of population structure inferred from core-genome (S) defined above, and one-hot encoded 3) matrix of SNP sites (SNP), where SNP_ij_ = 0 if strain *j* carries the ancestral allele at site *i*, and 1, 2, 3, 4, 5 if it contained A, T, C, G nucleotide or missing information, respectively; and 4) matrix of years of isolation (Y). We standardized each feature to have 0 mean and unit variance. Genes, strain clusters, and SNPs with identical indicator pattern were collapsed, so there are no duplicate rows in the G, S, or SNP matrices.

### Resistance prediction

We performed prediction using various combinations of input predictor matrices using resistance indicator as the output. We used 70% of the data for training the various models, and used the F1 score (harmonic mean of precision and recall) for resistance classification to evaluate them model on the remaining 30%. Four different models were used along with a baseline:

- Logistic regression with L_2_ regularization. We employed the “LogisticRegression” function in the Scikit-learn python package (www.scikit-learn.org) [36], with the “lbfgs” solver, and varied the regularization parameter strength from 0 to 1 with step size 0.01.

- Random forest classifier. We employed the “RandomForestClassifier” function in Scikit-learn and varied the number of trees in the forest (100, 300 and 600). When searching for the best split, we used both square root and binary logarithm selection of the number of features. We used bootstrap samples for building trees, out-of-bag samples to estimate accuracy, Gini impurity as the criterion for the information gain, and trained until all leaves were pure.

- Gradient boosted decision trees. We used the “GradientBoostingClassifier” implementation in Scikit-learn, with learning rate 0.1, and 100, 300 and 600 boosting stages. We used the same methods as for the random forests for choosing the number of features when selecting for the best split, and employed the deviance loss function. We limited the maximum depth of trees to 3, and the minimum number of samples required to split an internal or leaf node to 2 and 1, respectively. In order to assess the robustness of feature importance analysis, we repeated the optimization with 50 random seeds. As a measure for feature importance, we counted the number of times a feature used in optimization, as well as the average feature rank and importance across multiple replica.

- Deep neural networks. We employed the keras library in python (www.keras.io) to build fully connected deep neural networks. We tested two and four layer networks, with two output nodes corresponding to resistant and susceptible states, and 200, 300 and 400 nodes in each internal layer. We used Adam to train for 20 epochs, with batch size of 128, learning rate of 0.1, drop-out of 0, 0.1 or 0.3, and stopping when the validation set performance decreased. Due to the small training dataset size compared to the number of features, for ∼50% of runs the loss in the validation did not decrease by the end of training the network. We randomly partitioned the data into training (56%), validation (14%) and test (30%) sets, and trained models with different parameters on the training set, evaluated quality on the validation set, and final performance on the test set.

- Rule-based baseline. We compared our results with a rule-based method based on the detection of known resistance genes. To this end, we employed srst2 [29] and mapped short reads to the ResFinder database of known resistance genes in the srst2 package, using the cut-off of 60% for the length coverage.

## Results

Our data comprise 1936 samples that have been full genome sequenced, and phenotyped for resistance of 11 antibiotics. Resistance was distributed both within specific clades as well as emerging sporadically on divergent lineages (Figure S1), with an average frequency of 0.35 per drug (range: 0.15-0.63). This pattern is suggestive of both vertical and horizontal spread of resistance determinants. Genome sequences were processed to give gene and polymorphism presence information (1,390 core genes present in >99% of lineages, 90,261 genes present in one than one lineage, 1,432,145 variable sites in core genes), and 1,071 population structure features.

We used these predictors to test the ability of four machine learning models - logistic regression, random forests, gradient boosted decision trees, and deep neural networks - to predict antibiotic resistance. We varied model hyperparameters, as well as input data types, to establish the best-in-class predictors according to the F1 score for resistance (Methods). Gradient boosted decision trees performed best for predicting resistance of 11/11, and susceptibility of 10/11 drugs (Figure 1), with average precision of 0.93 and recall of 0.83 (Figure S2). Perhaps surprisingly, deep learning models that account for complex non-linear relations amongst features did not provide substantial improvement over the simpler logistic regression models, or random forests (Figure 1).

**Figure 1.**
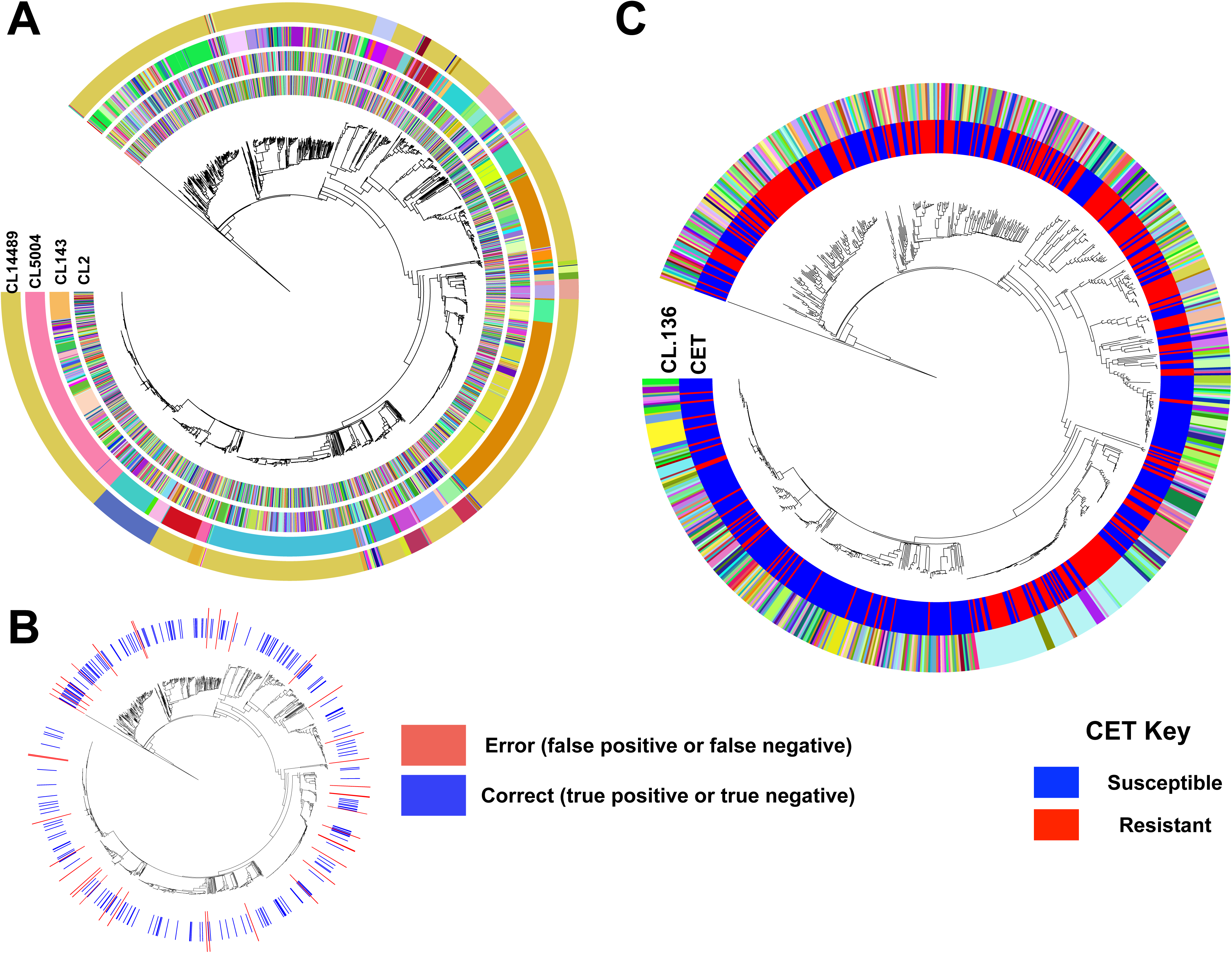
Prediction performance of the best tuned models. F1 score (harmonic mean of precision and recall; y-axis) for resistant (top panel) and susceptible (middle panel) phenotypes for four predictive models (red: gradient boosted decision trees; green: logistic regression; teal: random forests; purple: deep learning) across eleven antibiotics (x-axis). The best model of each class for every drug (x-axis) was identified based on the F1 score for resistance and employed a number of possible combinations of gene presence, population structure, and year of isolation (lower panel; black: feature used; white: feature not used).

Knowledge of what features that aid prediction will help prioritize data collection and diagnostic efforts. The gene presence and absence predictor (G) was used in all the best predictive models for each of the considered methods (Figure 1; lower panel). This is not surprising, given multiple known resistance mechanisms driven by accessory gene content, e.g. for beta-lactams and aminoglycosides. Population structure information (S) and year of isolation data (Y) were also frequently beneficial (used in 26 and 34 out of 44 best models, respectively). Adding gene presence to population structure features improved F1 score by 0.12 on average (Figure S3). In contrast, once gene presence had been accounted for, there was limited performance gain when including population structure features (Figure S3). This suggests that accessory gene content already contains information about population structure, which reflects the pattern of polymorphisms in the core genome. Indeed, core genome distance and accessory gene difference matrices are not independent (p < 0.01, Mantel test), which is likely explained by accessory genes acquired by clade ancestors, followed by limited turnover.

Next, we asked which individual features are most frequently utilized. We measured feature importance as the number of times it was used for gradient boosted decision trees, the best performing method, across 50 random fitting replicates on fixed training date (Figure S4). Overall, only an average 3% of input features (653 of 17198) were used at all in prediction across different drugs. In general, known resistance genes were identified as the most important, and were most frequently used features for predicting resistance to beta-lactams and aminoglycosides, e.g. *blaOXA-2, blaTEM-1, blaCTX-M-15*, and extra copies of *ampC* and *phnP* efflux pump genes (Supplemental Table S2). For example, the known beta-lactamase *bla-CTX-M* gene ranked first in all models for predicting resistance to beta-lactam ceftazidime, which followed by some genes with unknown function and *ampC* (Figure S4A). Perhaps surprisingly, we found that while the year of isolation was a dispensable feature for nearly all drugs, it was deemed important for ampicillin resistance prediction (Figure S4B). This was explained by the temporal distribution of the data, where all the strains collected in 2015 were resistant. These findings demonstrate that although known resistance genes were most predictive, other features, i.e. population structure and year of isolation, may be reproducibly used for prediction as well (Figure S4B). Nevertheless, it is clear that the inclusion of some features, such as collection year, reflects bias in the training data rather than biological importance.

Population structure information was often selected for use in the best performing models, and therefore useful for prediction (Figure 1). Indeed, training only on population structure produced an average F1 score of 0.73 (range: 0.23-0.92), and this performance could not be achieved with randomized phenotypes (Figure S5). Population structure features capture both recently diverged and deep clades (Figure 2A), and features included in the models were not limited to a single lineage or common depth (for example Figure 2B). As an example, the CL136 feature, which is the most important identified feature, distinguishes clusters by positing a maximum pairwise sequence distance between isolates of 136 nucleotides. Cluster membership at this level of similarity informs of resistance, as 85% of clusters with at least two strains contained either only resistant or only susceptible strains (Figure 2C). In these cases, resistance status of an ancestral strain of the clades was likely retained in descendants and did not change due to horizontal gene transfer, mutation, or sporadic gene loss. Altogether, the results show that predictive models can utilize genetic relatedness and population structure for predicting resistance, as has been observed in traditional eukaryotic genetics [8].

**Figure 2.**
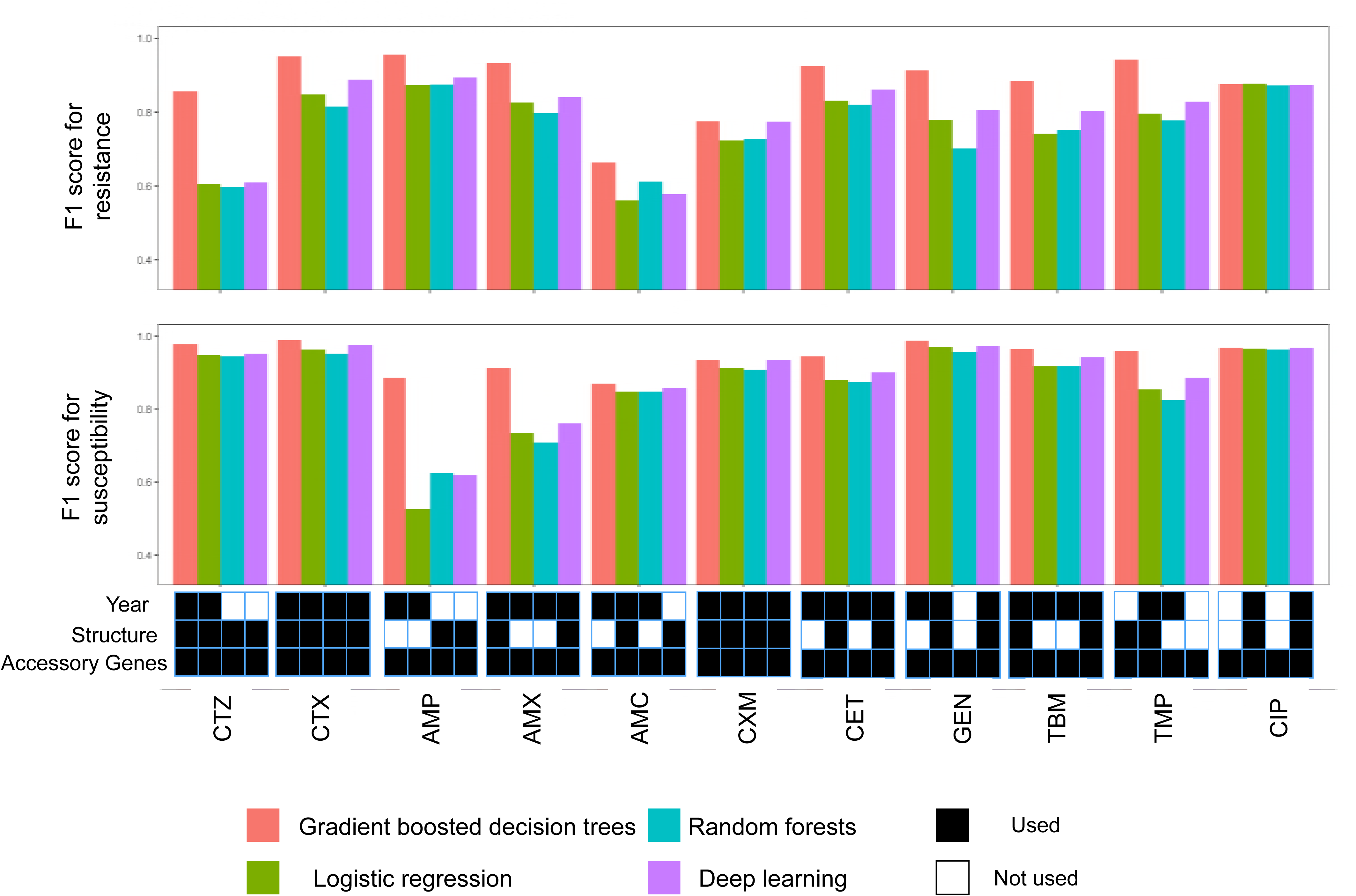
Population structure and phenotypic distribution of the input data. A) Phylogenetic distribution of clusters identified in the population for SNP distance cut-off values of 2, 143, 5054 and 14489 (outer circles) relative to the phylogenetic tree. B) Phylogenetic distribution of correct calls (true positives, true negatives) and errors (false positives, false negatives) when predicting cephalothin (CET) resistance with the best performing gradient boosted model. The F1 score for resistance was 0.81. C) Phylogenetic distribution of the most important identified population structure feature, clustering with SNP cut-off of 136 (outer ring), compared with the phylogenetic distribution of resistance phenotype (inner ring; blue: susceptible; red: resistant).

While the major mechanism for evolving antibiotic resistance is gene acquisition, mutations on chromosomes may also play a role, and therefore aid prediction. We thus next included single nucleotide polymorphism (SNP) data for gradient boosted decision trees, re-fitted hyperparameters, and evaluated on held out data. Predictive performance improved for four of the 11 antibiotics (Figure S3B). As anticipated, the largest improvement of 7.6% occurred for ciprofloxacin, resistance against which is known to involve chromosomal mutations [30, 31]. Accordingly, the three most important identified features were variants in chromosomal quinolone-resistance-determining regions of the genes encoding DNA gyrase (*gyrA*), and topoisomerase IV *parC*. For other antibiotics, the addition of SNP data either did not greatly improve or worsened prediction performance (Figure S3B, Discussion).

A possible limitation for applying machine learning methods to detect antibiotic resistance is the unavoidably small number of samples (1,936 in this study) compared to the number of features (∼18,270 in this study after collapsing the fully correlated features). To better understand how this imbalance impacts performance, we simulated data from different sample sizes using a range of penetrances for a single resistance determinant. As anticipated, the performance of gradient boosted decision trees dropped when the penetrance of the resistance determinant decreased (Figure S6). However, there was no reduction in F1 score upon decreasing the population size, even when using only 130 strains. Overall, these findings suggest that the large number of features relative to sample size does not impact model performance for high frequency causal genes.

The current clinical standards employ rule-based models to predict resistance from a small number of known determinants. We used *srst2* [29] to identify known resistance genes for cephalosporins, penicillins, aminoglycosides and trimethoprim (Table 2), and used this information to better understand prediction errors. The best model’s false positive resistance calls for different antibiotics contained 2 or 33 isolates that carried known resistance genes (beta-lactams, aminoglycoside modifying enzymes and *dfr* genes), but were annotated as susceptible. Manual inspection confirmed that all of these genes were fully covered by sequence data, and almost identical to the known resistance genes. Moreover, for different antibiotics, one to nine false negative resistance calls (36 total) did not contain a known causal determinant. Similarly, two to nine resistant strains (51 total) were correctly marked as resistant by our model, but contained no known resistance gene (Table 2). These discrepancies may be explained by either resistance testing error, genomic sequence quality, or unknown mechanisms for resistance. As neither approach was perfect, predictive models in combination with rule-based methods may help identify cases that necessitate further analysis or repeating the susceptibility tests, ultimately leading to improved diagnostics and novel mechanisms.

**Table 1.**
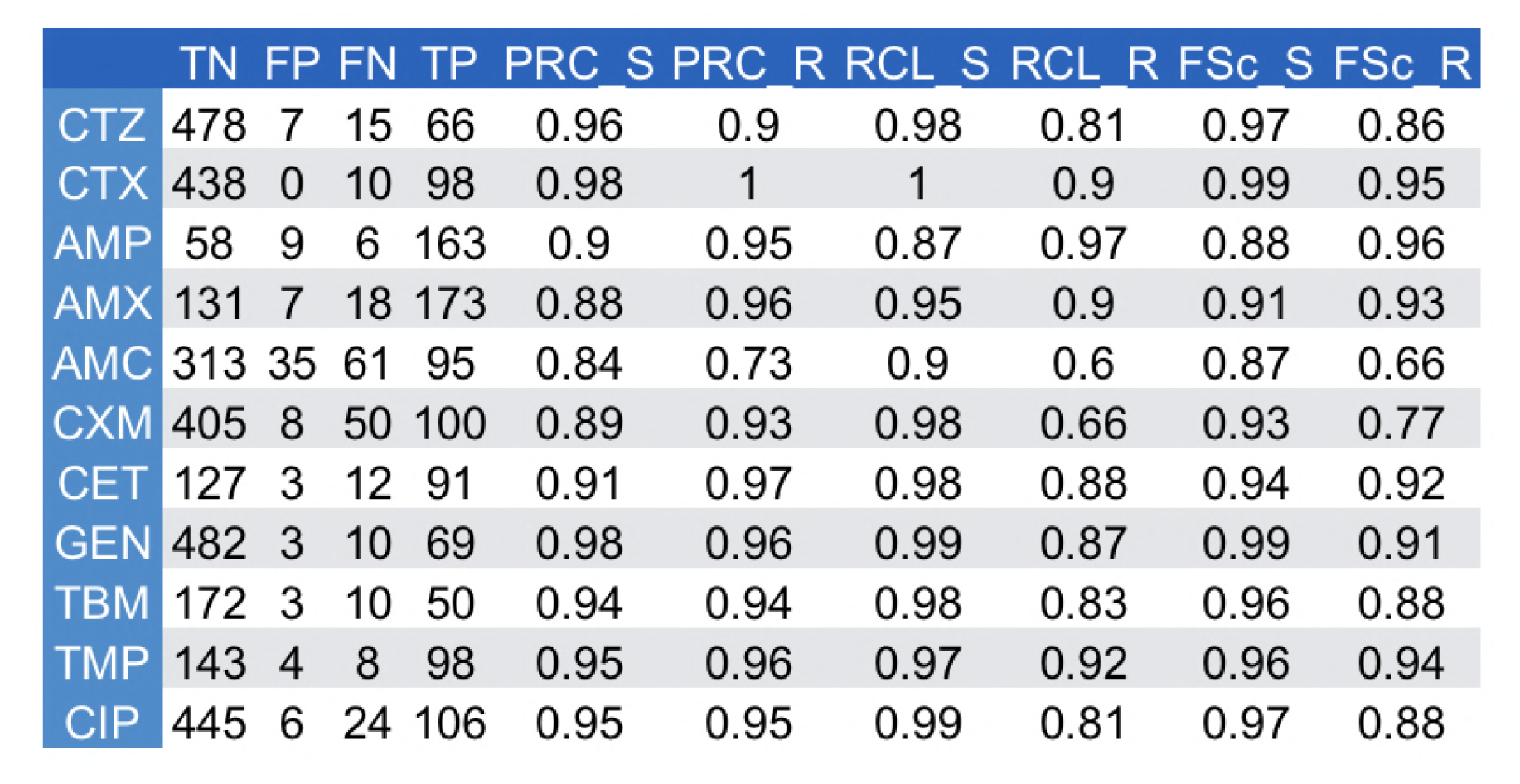
Prediction metrics on held out data for the best performing gradient boosted decision trees model. TN: true negatives, FN: false negatives, FP: false positives, TP: true positives, PRC: precision, RCL: recall, S: susceptibility, R: resistance.

**Table 2.** Comparison of prediction results with a rule-based model, srst2. True positives (TP), false positives (FP) and false negatives (FN) from Table 1. Resistant genes were identified by srst2 using the Resfinder database. Since ciprofloxacin resistance is caused by chromosomal mutations, we have not included this antibiotic in the table.

| Drug | TP without resistant gene | FN without resistant gene | FP with resistant gene |
| --- | --- | --- | --- |
| CTZ | 3/66 | 1/15 | 7/7 |
| CTX | 2/98 | 4/10 | - |
| AMP | 9/163 | 1/6 | 2/9 |
| AMX | 8/173 | 0/18 | 5/7 |
| AMC | 0/95 | 1/61 | 33/35 |
| CXM | 3/100 | 4/50 | 6/8 |
| CET | 6/91 | 7/12 | 3/3 |
| GEN | 7/69 | 9/10 | 2/3 |
| TBM | 7/50 | 2/10 | 3/3 |
| TMP | 6/98 | 7/8 | 1/4 |

## Discussion

We examined the ability of four different machine learning methods to predict antibiotic resistance from genomic information in *E.coli*, without making assumptions about the underlying genetic mechanisms. Our tests revealed that accessory genome data is needed for high accuracy in general, but that population structure information can also aid prediction.

Our input dataset was diverse. The collection comprised seven sequence types and strains from 15 consecutive years across a range of geographical locations. The majority of isolates (1509 of 1936 samples) were from a nationwide study across hospitals in the United Kingdom and Ireland [15], and associated with bacteremia. This geographical bias is not expected to affect the performance of the model on a new clinical dataset, since *E. coli* sequence types (e.g. ST131 clone [32]) are circulated across hospitals worldwide. However, as isolates from potential reservoirs, including hospital sewage and wastewater treatment plants, were underrepresented in training data (99 of 1936 samples), we cannot conclusively assess how well the trained models detect resistance in samples from these sources. More data are needed to develop robust models across the entire species range, especially if resistance mechanisms differ in the various niches.

The phenotype data was binary - each isolate was deemed either resistant or not to a compound. It is clear that this is an oversimplification of reality, as substantial variation hides within both categories. As the resistance phenotype is directly exposed to selection, it will influence how quickly it spreads within and between patients, as well as in bacterial populations at large. To predict treatment outcomes, correctly design interventions and allocate societal resources, it will therefore be important to be able to accurately predict resistance quantitatively as well. This requires non-binary resistance data, acquired at high accuracy and throughput.

Our findings confirm the utility of ensemble methods, and in particular boosting models, for predicting antibiotic resistance. While deep learning models are able to capture higher order interactions between features, and therefore often outperform simpler alternatives [37], they did not provide additional advantage here. Tree-based methods are often used as an intermediate between simple models that treat features independently, like logistic regression, and more complex, but poorly interpretable models. Indeed, random forest readouts can be analysed for feature importance as we have done here, and even detecting genetic interactions (e.g. [33, 34]). However, as for association methods [28, 35], the true impact of genetic features is confounded by their phylogenetic distribution and population structure. Therefore, approaches to distinguish causal resistance genes from all correlated markers require additional experimental study.

Recent reports have confirmed the strength of tree-based methods for predicting clinical attributes. For example, Wheeler *et al*. used random forests to predict invasiveness of *Salmonella enterica* lineages [14]. In another study, a tree ensemble was trained with boosting to predict the minimum inhibitory concentration from DNA k-mers for a large-scale *Klebsiella pneumoniae* panel [10], but the value of using core genome compared to accessory genes was not investigated. In general, including variant data or k-mers in the model greatly increases the number of features. However, adding the ∼1 million additional single nucleotide features to ∼20,000 others did not improve the results for most drugs in our dataset. This suggests that only pan-genome data could be used in early screenings for resistance, and including nucleotide-level information will be more beneficial in a limited form, once causality is established for a broader range of SNPs.

Many factors can affect the performance of a predictive model. For instance, we noted that amongst the antibiotics, the results were worst for amoxicillin-clavulanate. This might be due to a beta-lactamase inhibitor, which dominates the impact of a resistance gene, causing susceptibility, and resulting in a wrong prediction. The genome-based prediction also cannot account for non-genomic resistance mechanisms, such as high expression level of resistance genes. Consequently, future models should assess the value of even broader data for accurate prediction, ranging from transcriptome and proteome to other clinical and epidemiological data, such as cross-resistance and history of antibiotic therapy. Integrating these information sources from large isolate panels into a single predictive framework will lead to a rational basis for decision-making in public health to reduce cost of diagnoses and treatments.

## Author Contributions

DM and LP designed the study framework. DM developed and implemented the model. LP, VM and JW supervised the project. MP and AF generated data. All authors read and approved the final manuscript. The authors have declared that no competing interests exist.

## Funding information

This work was partially funded by a grant from the Centre for Antibiotic Resistance Research (CARe) at the University of Gothenburg to AF and grant number 2016-06503 from the Joint Programming Initiative on Antimicrobial Resistance (JPIAMR) to JW and AF. This work was in part supported by the Academy of Finland (grant 313270 to VM). LP was supported by Wellcome, and Estonian Research Council (IUT34-4). DM was supported by the Joint Programming Initiative on Antimicrobial Resistance (JPIAMR) via MRC grant MR/R004501/1. The funders had no role in study design, data collection and analysis, decision to publish, or preparation of the manuscript.

### Acknowledgements

Christina Åhrén, Nahid Karami, Carl-Fredrik Flach, Ed Moore (Culture Collection University of Gothenburg, CCUG), Jan Michiels and Marco Galardini are gratefully acknowledged for providing strains. Daniel Jaén Luchoro, Fabrice E. Graf, Owens Uwangue and Viktor Garellick are gratefully acknowledged for technical assistance and helpful discussions.

## Supplemental Tables

**Supplemental Table S1: list of isolates with associated metadata and accession numbers in the European Nucleotide Archive (ENA).**

**Supplemental Table S2: List of important accessory genes and their functions for feature importance analysis with the best performing gradient boosted decision trees models shown in Figure S4. The importance metrics include the number of runs (total 50 runs), in which the feature was used during model optimization and the average ranking and importance for the feature in these runs.**

